# Identifying Parkinson’s disease and parkinsonism cases using routinely-collected healthcare data: a systematic review

**DOI:** 10.1101/331652

**Authors:** Zoe Harding, Tim Wilkinson, Anna Stevenson, Sophie Horrocks, Amanda Ly, Christian Schnier, David P Breen, Kristiina Rannikmäe, Cathie LM Sudlow, on behalf of Dementias Platform UK

## Abstract

**Background:** Population-based, prospective studies can provide important insights into Parkinson’s disease (PD) and other parkinsonian disorders. Participant follow-up in such studies is often achieved through linkage to routinely-collected healthcare datasets. We systematically reviewed the published literature on the accuracy of these datasets for this purpose.

**Methods:** We searched four electronic databases for published studies that compared PD and parkinsonism cases identified using routinely-collected data to a reference standard. We extracted study characteristics and two accuracy measures: positive predictive value (PPV) and/or sensitivity.

**Results:** We identified 18 articles, resulting in 27 measures of PPV and 14 of sensitivity. For PD, PPVs ranged from 56-90% in hospital datasets, 53-87% in prescription datasets, 81-90% in primary care datasets and was 67% in mortality datasets. Combining diagnostic and medication codes increased PPV. For parkinsonism, PPVs ranged from 36-88% in hospital datasets, 40-74% in prescription datasets, and was 94% in mortality datasets. Sensitivities ranged from 15-73% in single datasets for PD and 43-63% in single datasets for parkinsonism.

**Conclusions:** In many settings, routinely-collected datasets generate good PPVs and reasonable sensitivities for identifying PD and parkinsonism cases. Further research is warranted to investigate primary care and medication datasets, and to develop algorithms that balance a high PPV with acceptable sensitivity.

## Introduction

Despite well-established pathological features, the aetiologies of Parkinson’s Disease (PD) and other parkinsonian conditions remain poorly understood and disease-modifying treatments have proved elusive. Large, prospective, population-based cohort studies with biosample collections (e.g., UK Biobank, German National Cohort, US Precision Medicine Initiative) provide a robust methodological framework with statistical power to investigate the complex interplay between genetic, environmental and lifestyle factors in the aetiology and natural history of neurological disorders such as PD and other parkinsonian disorders[1–3].

Linkage to routinely-collected healthcare data – which are administrative datasets collected primarily for healthcare purposes rather than to address specific research questions[4] –provides an efficient means of long term follow-up in order to identify large numbers of incident cases in such studies[1]. Furthermore, participant linkage to such datasets can be used in randomised controlled trials as a cost-effective and comprehensive method of follow-up for disease outcomes[5]. These data are coded using systems such as the International Classification of Diseases (ICD)[6], the Systematized Nomenclature of Medicine – Clinical Terms (SNOMED-CT) system[7], and the UK primary care Read system[8].

Before such datasets can be used to identify PD and parkinsonism cases in prospective studies, their accuracy must be determined. Important measures are the positive predictive value (PPV, the proportion of those coded positive that are true disease cases) and sensitivity (the proportion of true disease cases that are coded positive). Specificity and negative predictive value (NPV) are less relevant as specificity will be high when precise diagnostic codes are used and NPV, which is related to disease prevalence, will be high in population-based studies where most individuals do not develop the disease of interest.

We systematically reviewed published studies evaluating the accuracy of routinely-collected healthcare data for identifying PD and parkinsonism cases.

## Methods

### Study Protocol

We prospectively published the protocol for this systematic review (www.crd.york.ac.uk/PROSPERO, number: 2016:CRD42016033715)[9].

### Search Strategy and Eligibility Criteria

We searched the electronic databases MEDLINE (Ovid), EMBASE (Ovid), CENTRAL (Cochrane Library) and Web of Science (Thomson Reuters) for articles published in any language between 01.01.1990 and 23.06.2017 that compared codes for PD or parkinsonism from routinely-collected healthcare data to a clinical expert-derived reference standard (see Supplementary File S1 for search strategy). Studies had to provide either a PPV and/or a sensitivity estimate, or sufficient raw data to calculate these. Where articles assessed more than one dataset or evaluated both PPV and sensitivity, we included these as separate studies. Hereafter we will refer to published papers as ‘articles’ and these separate analyses as ‘studies’. We chose the date limits based on our judgement that accuracy estimates from studies published prior to 1990 would have limited current applicability. We also screened bibliographies of included studies and relevant review papers to identify additional publications. Studies had to have ≥10 coded cases, due to the limited precision of studies below this size. Studies reporting sensitivity values had to be population-based (i.e. community-based as opposed to hospital-based) with comprehensive attempts to detect all disease cases. Where multiple studies investigated overlapping populations, we included the study with the larger population size.

### Study Selection

Two authors (AS and SH) independently screened all titles and abstracts generated by the search, and reviewed full text articles of all potentially eligible studies to determine if the inclusion criteria were met. In the case of disagreement or uncertainty, we reached a consensus through discussion and, where necessary, involvement of a senior third author (CLMS).

### Data Extraction

Using a standardized form, two authors (TW and ZH) independently extracted the following data from each study: first author, year of publication, time period during which coded data were collected, country of study, study population, study size (defined as the total number of code positive cases for PPV [true positives plus false positives] and the total number of true positives for sensitivity [true positives and false negatives]), type of routine data used (e.g., hospital admissions, mortality or primary care), coding system and version used, specific codes used to identify cases, diagnostic coding position (e.g. primary or secondary position), parkinsonian subtypes investigated, and the method used to make the reference standard diagnosis.

We recorded the reported PPV and/or sensitivity estimates, as well as any corresponding raw data. After discussion, any remaining queries were resolved with a senior third author (CLMS). When necessary, we contacted study authors to request additional information.

### Quality Assessment

We adapted the Quality Assessment of Diagnostic Accuracy Studies 2 (QUADAS-2)[10] tool to evaluate the risk of bias in the estimates of accuracy and any concerns about the applicability of each article to our specific research question (Supplementary Table S2). Two authors (TW and ZH) independently assigned quality ratings, with any discrepancies resolved through discussion. We performed this evaluation in the context of our specific review question and not as an indication of the overall quality of the articles.

### Statistical Analysis/Data Synthesis

We tabulated the extracted data, and calculated 95% confidence intervals for the accuracy measures from the raw data using the Clopper-Pearson (exact) method. Due to substantial heterogeneity in study settings and methodologies, we did not perform a meta-analysis, as we considered any summary estimate to be potentially misleading. Instead, we assessed the full range of results in the context of study methodologies, populations and specific data sources. We also reported any within-study comparisons in which a single variable was changed to examine its effect on PPV or sensitivity. We performed analyses using the statistical software StatsDirect3.

## Results

### Study Characteristics

18 published articles fulfilled our inclusion criteria[11–28]. A flow diagram of the study selection process is shown in Fig 1. We obtained key additional information from the authors of two studies[20,24]. Of the 18 included articles, 13 reported PPV[11,13–24], four reported sensitivity[25–28] and one reported both[12]. Four articles contained more than one study[11–13,17]. One of these consisted of multiple sub-studies, using different methods to evaluate datasets across several countries, so we included these as six separate studies[13]. In total, there were 27 measures of PPV and 14 of sensitivity. Study characteristics are summarised in Tables 1 and 2 respectively.

**Fig 1:**
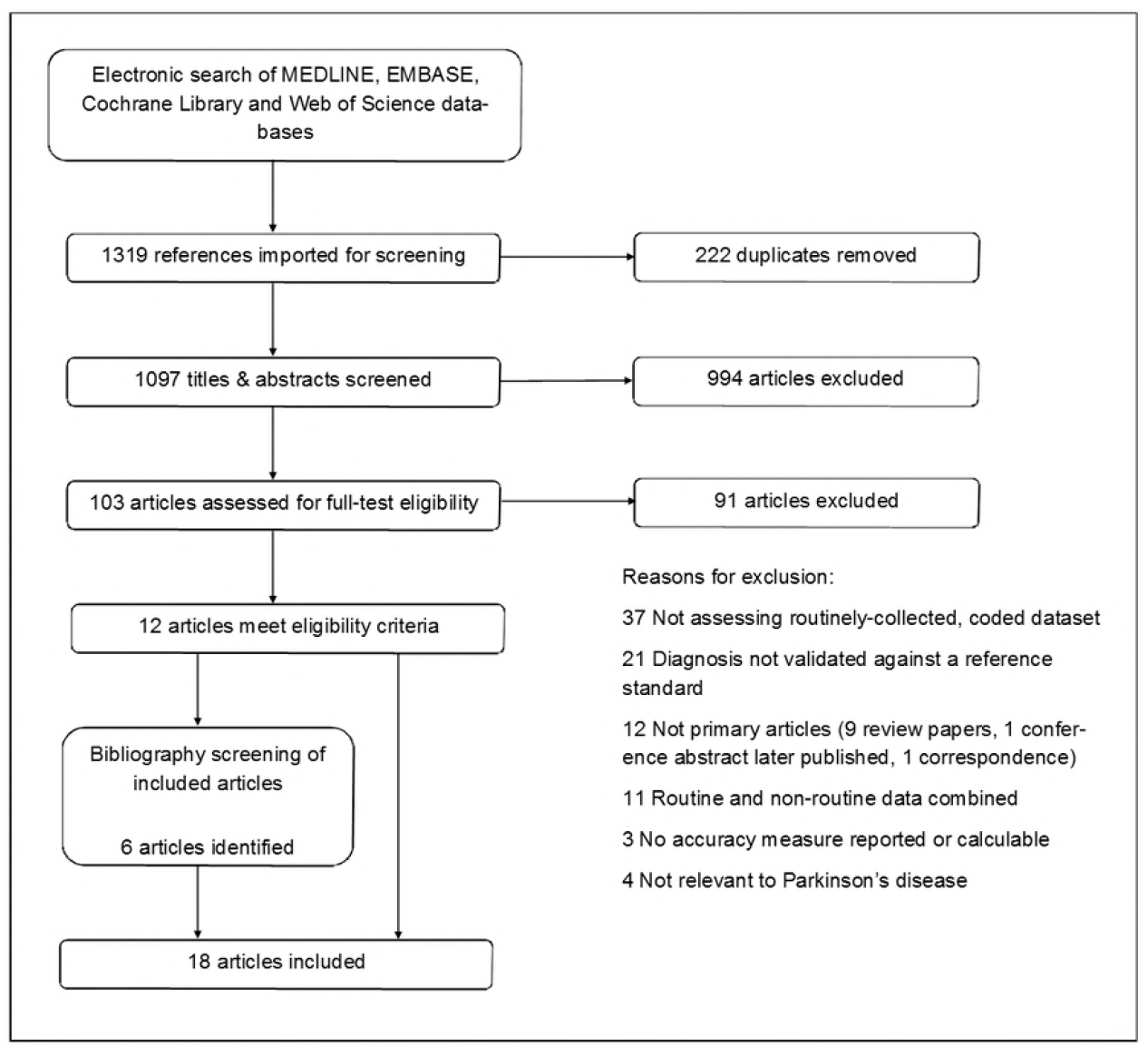
PRISMA Flow Diagram.

**Table 1.**
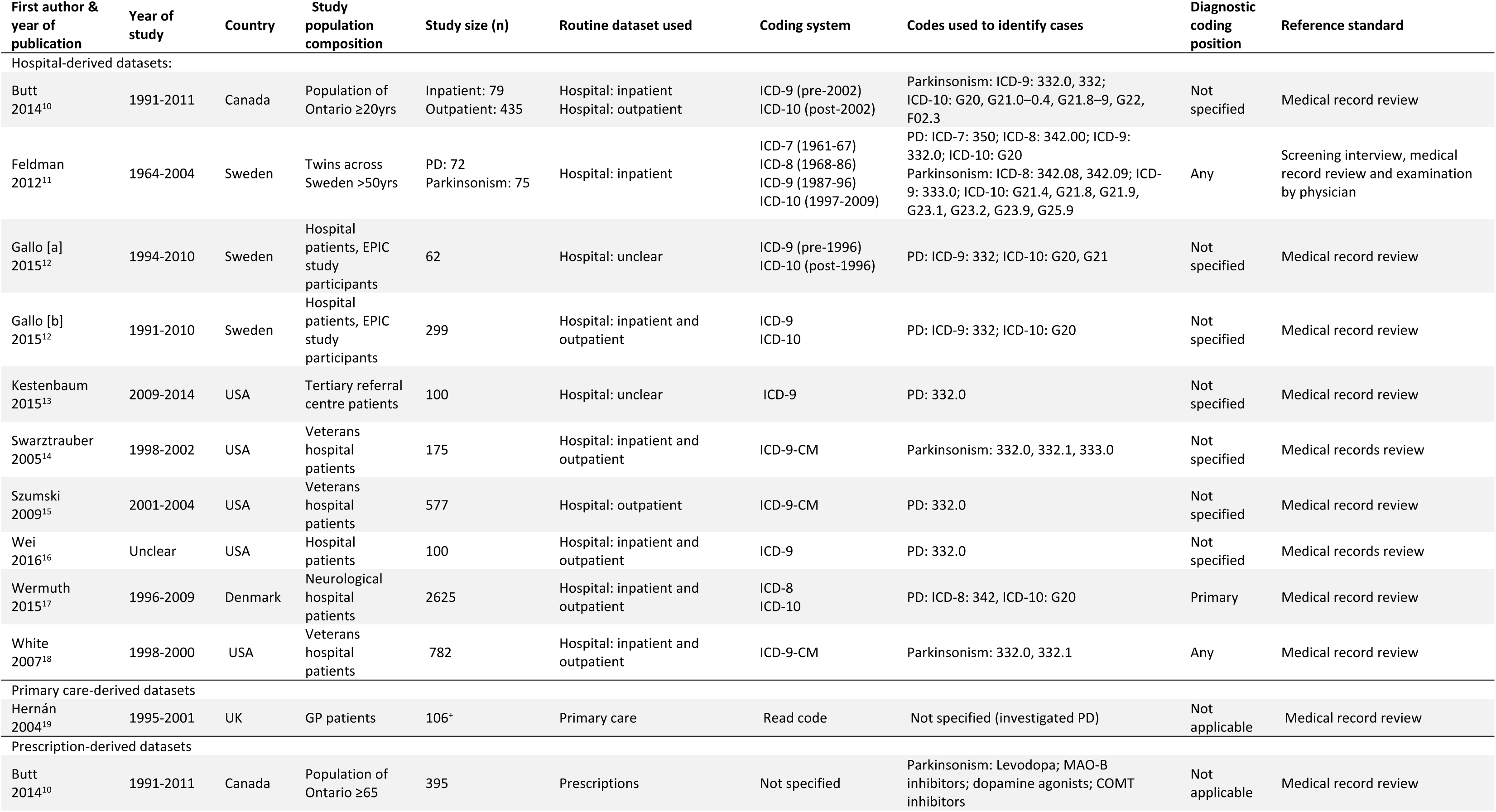
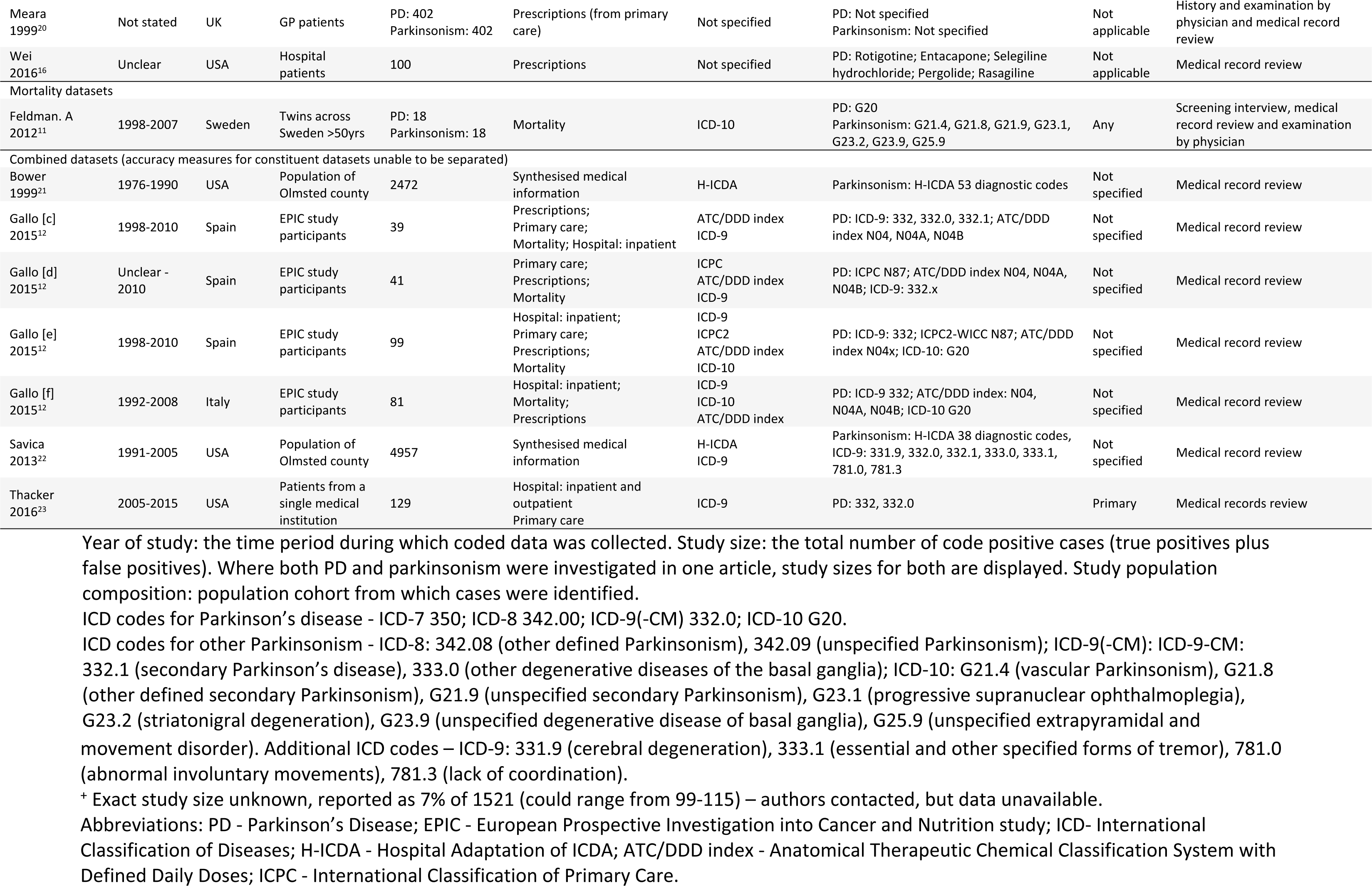
Characteristics of studies reporting positive predictive value, stratified by dataset type.

| First author & year of publication | Year of study | Country | Study population composition | Study size (n) | Routine dataset used | Coding system | Codes used to identify cases | Diagnostic coding position | Reference standard |
| --- | --- | --- | --- | --- | --- | --- | --- | --- | --- |
| Hospital-derived datasets: |  |  |  |  |  |  |  |  |  |
| Butt 2014<sup>10</sup> | 1991-2011 | Canada | Population of Ontario ≥20yrs | Inpatient: 79<br>Outpatient: 435 | Hospital: inpatient<br>Hospital: outpatient | ICD-9 (pre-2002)<br>ICD-10 (post-2002) | Parkinsonism: ICD-9: 332.0, 332;<br>ICD-10: G20, G21.0–0.4, G21.8–9, G22, F02.3 | Not specified | Medical record review |
| Feldman 2012<sup>11</sup> | 1964-2004 | Sweden | Twins across Sweden >50yrs | PD: 72<br>Parkinsonism: 75 | Hospital: inpatient | ICD-7 (1961-67)<br>ICD-8 (1968-86)<br>ICD-9 (1987-96)<br>ICD-10 (1997-2009) | PD: ICD-7: 350; ICD-8: 342.00; ICD-9: 332.0; ICD-10: G20<br>Parkinsonism: ICD-8: 342.08, 342.09; ICD-9: 333.0; ICD-10: G21.4, G21.8, G21.9, G23.1, G23.2, G23.9, G25.9 | Any | Screening interview, medical record review and examination by physician |
| Gallo [a] 2015<sup>12</sup> | 1994-2010 | Sweden | Hospital patients, EPIC study participants | 62 | Hospital: unclear | ICD-9 (pre-1996)<br>ICD-10 (post-1996) | PD: ICD-9: 332; ICD-10: G20, G21 | Not specified | Medical record review |
| Gallo [b] 2015<sup>12</sup> | 1991-2010 | Sweden | Hospital patients, EPIC study participants | 299 | Hospital: inpatient and outpatient | ICD-9<br>ICD-10 | PD: ICD-9: 332; ICD-10: G20 | Not specified | Medical record review |
| Kestenbaum 2015<sup>13</sup> | 2009-2014 | USA | Tertiary referral centre patients | 100 | Hospital: unclear | ICD-9 | PD: 332.0 | Not specified | Medical record review |
| Swarztrauber 2005<sup>14</sup> | 1998-2002 | USA | Veterans hospital patients | 175 | Hospital: inpatient and outpatient | ICD-9-CM | Parkinsonism: 332.0, 332.1, 333.0 | Not specified | Medical records review |
| Szumski 2009<sup>15</sup> | 2001-2004 | USA | Veterans hospital patients | 577 | Hospital: outpatient | ICD-9-CM | PD: 332.0 | Not specified | Medical record review |
| Wei 2016<sup>16</sup> | Unclear | USA | Hospital patients | 100 | Hospital: inpatient and outpatient | ICD-9 | PD: 332.0 | Not specified | Medical records review |
| Wermuth 2015<sup>17</sup> | 1996-2009 | Denmark | Neurological hospital patients | 2625 | Hospital: inpatient and outpatient | ICD-8<br>ICD-10 | PD: ICD-8: 342, ICD-10: G20 | Primary | Medical record review |
| White 2007<sup>18</sup> | 1998-2000 | USA | Veterans hospital patients | 782 | Hospital: inpatient and outpatient | ICD-9-CM | Parkinsonism: 332.0, 332.1 | Any | Medical record review |
| Primary care-derived datasets |  |  |  |  |  |  |  |  |  |
| Hernán 2004<sup>19</sup> | 1995-2001 | UK | GP patients | 106* | Primary care | Read code | Not specified (investigated PD) | Not applicable | Medical record review |
| Prescription-derived datasets |  |  |  |  |  |  |  |  |  |
| Butt 2014<sup>10</sup> | 1991-2011 | Canada | Population of Ontario ≥65 | 395 | Prescriptions | Not specified | Parkinsonism: Levodopa; MAO-B inhibitors; dopamine agonists; COMT inhibitors | Not applicable | Medical record review |
| Meara 1999<sup>20</sup> | Not stated | UK | GP patients | PD: 402<br>Parkinsonism: 402 | Prescriptions (from primary care) | Not specified | PD: Not specified<br>Parkinsonism: Not specified | Not applicable | History and examination by physician and medical record review |
| Wei 2016<sup>16</sup> | Unclear | USA | Hospital patients | 100 | Prescriptions | Not specified | PD: Rotigotine; Entacapone; Selegiline hydrochloride; Pergolide; Rasagiline | Not applicable | Medical record review |
| Mortality datasets |  |  |  |  |  |  |  |  |  |
| Feldman. A 2012<sup>11</sup> | 1998-2007 | Sweden | Twins across Sweden >50yrs | PD: 18<br>Parkinsonism: 18 | Mortality | ICD-10 | PD: G20<br>Parkinsonism: G21.4, G21.8, G21.9, G23.1, G23.2, G23.9, G25.9 | Any | Screening interview, medical record review and examination by physician |
| Combined datasets (accuracy measures for constituent datasets unable to be separated) |  |  |  |  |  |  |  |  |  |
| Bower 1999<sup>21</sup> | 1976-1990 | USA | Population of Olmsted county | 2472 | Synthesised medical information | H-ICDA | Parkinsonism: H-ICDA 53 diagnostic codes | Not specified | Medical record review |
| Gallo [c] 2015<sup>12</sup> | 1998-2010 | Spain | EPIC study participants | 39 | Prescriptions;<br>Primary care;<br>Mortality; Hospital: inpatient | ATC/DDD index<br>ICD-9 | PD: ICD-9: 332, 332.0, 332.1; ATC/DDD index N04, N04A, N04B | Not specified | Medical record review |
| Gallo [d] 2015<sup>12</sup> | Unclear - 2010 | Spain | EPIC study participants | 41 | Primary care;<br>Prescriptions;<br>Mortality | ICPC<br>ATC/DDD index<br>ICD-9 | PD: ICPC N87; ATC/DDD index N04, N04A, N04B; ICD-9: 332.x | Not specified | Medical record review |
| Gallo [e] 2015<sup>12</sup> | 1998-2010 | Spain | EPIC study participants | 99 | Hospital: inpatient;<br>Primary care;<br>Prescriptions;<br>Mortality | ICD-9<br>ICPC2<br>ATC/DDD index<br>ICD-10 | PD: ICD-9: 332; ICPC2-WICC N87; ATC/DDD index N04x; ICD-10: G20 | Not specified | Medical record review |
| Gallo [f] 2015<sup>12</sup> | 1992-2008 | Italy | EPIC study participants | 81 | Hospital: inpatient;<br>Mortality;<br>Prescriptions | ICD-9<br>ICD-10<br>ATC/DDD index | PD: ICD-9 332; ATC/DDD index: N04, N04A, N04B; ICD-10 G20 | Not specified | Medical record review |
| Savica 2013<sup>22</sup> | 1991-2005 | USA | Population of Olmsted county | 4957 | Synthesised medical information | H-ICDA<br>ICD-9 | Parkinsonism: H-ICDA 38 diagnostic codes, ICD-9: 331.9, 332.0, 332.1, 333.0, 333.1, 781.0, 781.3 | Not specified | Medical record review |
| Thacker 2016<sup>23</sup> | 2005-2015 | USA | Patients from a single medical institution | 129 | Hospital: inpatient and outpatient<br>Primary care | ICD-9 | PD: 332, 332.0 | Primary | Medical records review |

Year of study: the time period during which coded data was collected. Study size: the total number of code positive cases (true positives plus false positives). Where both PD and parkinsonism were investigated in one article, study sizes for both are displayed. Study population composition: population cohort from which cases were identified. ICD codes for Parkinson's disease - ICD-7 350; ICD-8 342.00; ICD-9(-CM) 332.0; ICD-10 G20. ICD codes for other Parkinsonism - ICD-8: 342.08 (other defined Parkinsonism), 342.09 (unspecified Parkinsonism); ICD-9(-CM): ICD-9-CM: 332.1 (secondary Parkinson's disease), 333.0 (other degenerative diseases of the basal ganglia); ICD-10: G21.4 (vascular Parkinsonism), G21.8 (other defined secondary Parkinsonism), G21.9 (unspecified secondary Parkinsonism), G23.1 (progressive supranuclear ophthalmoplegia), G23.2 (striatonigral degeneration), G23.9 (unspecified degenerative disease of basal ganglia), G25.9 (unspecified extrapyramidal and
movement disorder). Additional ICD codes – ICD-9: 331.9 (cerebral degeneration), 333.1 (essential and other specified forms of tremor), 781.0 (abnormal involuntary movements), 781.3 (lack of coordination).
\* Exact study size unknown, reported as 7% of 1521 (could range from 99-115) – authors contacted, but data unavailable.
Abbreviations: PD - Parkinson's Disease; EPIC - European Prospective Investigation into Cancer and Nutrition study; ICD- International Classification of Diseases; H-ICDA - Hospital Adaptation of ICDA; ATC/DDD index - Anatomical Therapeutic Chemical Classification System with Defined Daily Doses; ICPC - International Classification of Primary Care.

**Table 2.** Characteristics of studies reporting sensitivity, stratified by dataset type.

| First author, year of publication | Year of study | Country | Study population composition | Study size (n) | Routine dataset used | Coding system | Codes used to identify cases | Diagnostic coding position | Reference standard |
| --- | --- | --- | --- | --- | --- | --- | --- | --- | --- |
| Mortality certificate-derived datasets: |  |  |  |  |  |  |  |  |  |
| Benito-León 2014<sup>24</sup> | 1994-2007 | Spain | Three communities near Madrid | 82 | Mortality | ICD-9 (pre 1999)<br>ICD-10 (post 1999) | Not specified (investigated PD) | Primary | Screening (in-person, telephone and mail questionnaire) and neurological examination |
| Beyer 2001<sup>25</sup> | 1993-1996 | Norway | County (Rogaland) | 84 | Mortality | ICD-9 or ICPC | Not specified (investigated PD) | Primary + Any | Semi-structured interview and a clinical examination |
| Fall 2003<sup>26</sup> | 1989-1998 | Sweden | Central district of Östergötland | 121 | Mortality | ICD-9 | Not specified (investigated PD) | Primary + Any | Examination and medical record review |
| Feldman 2012<sup>11</sup> | 1998-2008 | Sweden | Twins across Sweden >50yrs | Parkinsonism: 127<br>PD: 77 | Mortality | ICD-10 | PD: G20<br>Parkinsonism: G21.4, G21.8, G21.9, G23.1, G23.2, G23.9, G25.9 | Any | Screening interview, medical record review and examination |
| Williams-Gray 2013<sup>27</sup> | 2000-2012 | UK | County (Cambridgeshire) | 63 | Mortality | Not specified | Not specified (investigated PD) | Primary + Any | History and neurological examination |
| Hospital-derived datasets: |  |  |  |  |  |  |  |  |  |
| Feldman 2012<sup>11</sup> | 1964-2009 | Sweden | Twins across Sweden >50yrs | Parkinsonism: 194<br>PD: 132 | Hospital: inpatient | ICD-7 (1961-67)<br>ICD-8 (1968-86)<br>ICD-9 (1987-96)<br>ICD-10 (1997-2009) | PD: ICD-7: 350; ICD-8: 342.00; ICD-9: 332.0; ICD-10: G20<br>Parkinsonism: ICD-8: 342.08, 342.09; ICD-9: 333.0; ICD-10: G21.4, G21.8, G21.9, G23.1, G23.2, G23.9, G25.9 | Any | Screening interview, medical record review and examination |

Year of study: the time period during which coded data was collected. Study size: the total number of true positive according to the reference standard (true positives and false negatives). Where both PD and parkinsonism were investigated in one article, study sizes for both are displayed. Study population composition: population cohort from which cases were identified. ICD codes for Parkinson's disease - ICD-7 350; ICD-8 342.00; ICD-9 332.0; ICD-10 G20. ICD codes for other Parkinsonism - ICD-8: 342.08 (other defined Parkinsonism), 342.09 (unspecified Parkinsonism); ICD-9: 333.0 (other degenerative diseases of the basal ganglia); ICD-10: G21.4 (vascular Parkinsonism), G21.8 (other defined secondary Parkinsonism), G21.9 (unspecified secondary Parkinsonism), G23.1 (progressive supranuclear ophthalmoplegia), G23.2 (striatonigral degeneration), G23.9 (unspecified degenerative disease of basal ganglia), G25.9 (unspecified extrapyramidal and movement disorder)

Study size varied considerably, ranging from 39-4957. All 18 articles were based in high-income countries. Three were from the UK[20,21,28], six from mainland Europe[12,13,18,25–27], eight from the USA[14–17,19,22–24], and one from Canada[11]. There were 12 PPV estimates and two sensitivity estimates from hospital data[11–19], two PPV and 10 sensitivity estimates from mortality data[12,25–28], two PPV estimates from primary care data[20], four PPV estimates from prescription data[11,17,21] and seven PPV estimates and two sensitivity estimates from combining datasets from different sources[12,13,22–24]. There were no sensitivity estimates from primary care or prescription data.

PD was evaluated in 13 articles, with eight estimating PPV[13,14,16–18,20,21,24], four estimating sensitivity[25–28] and one estimating both[12]. Parkinsonism was evaluated by seven articles, of which six estimated PPV[11,15,19,21–23] and one assessed both PPV and sensitivity[12]. All of the parkinsonism articles combined PD with other causes of parkinsonism

The methods of reference standard used could be broadly divided into two categories: patient history and examination (majority of studies reporting sensitivity) and medical record review (majority of studies reporting PPV). In addition, where entire populations were under study, some studies incorporated a screening method (e.g., telephone interview) to identify potential cases[12,25].

Where reported, codes used to identify PD cases were consistent and appropriate to the ICD version used. However, the range of codes used to identify other parkinsonian conditions varied considerably, reflecting the broad range of pathologies that can lead to parkinsonism. Seven studies did not specify the exact codes used[17,20,21,25–28]. ICD versions used reflected the time period over which the studies were conducted. 19 studies used ICD-9 (or ICD-9-CM, a clinically modified version used in the USA, and identical to ICD-9 with respect to parkinsonian diagnoses)[11–17,19,23–27], 11 used ICD-10[11–13,18,25], three used ICD-8[12,18], and two used ICD-7[12]. One of the primary care studies used Read-coded data[20]. Four studies, including the three that evaluated prescription data, did not specify the coding system used[11,17,21,28].

The diagnostic coding position assessed also varied. Three studies assessed primary diagnoses alone[18,24,25], eight used any diagnostic position[12,19,26–28], while 13 did not specify the coding position[11,13–17,22,23]. Diagnostic position was not applicable in the studies of primary care and prescription data due to the nature of these datasets[11,17,20,21].

### Quality Assessment

Only two articles were judged to be of low risk of bias or applicability concerns in the QUADAS-2 assessment[11,12] (Supplementary Table S3). The commonest concerns were: selection bias, lack of reporting of the codes used to identify disease cases, insufficiently rigorous reference standards, inappropriate inclusions and exclusions, or patients being lost to follow-up.

### Positive predictive value

For PD, there were 17 PPV estimates in total (Fig 2)[12–14,16–18,20,21,24]. These comprised seven PPV estimates of hospital data alone[12–14,16–18], one of mortality data alone[12], two for prescription data alone[17,21], one of primary care data alone[20], one of prescription data and primary care data in combination[20], and five of datasets used in combination[13,24]. PPVs ranged from 36-90% across all studies. Nine of the 17 estimates were >75%. The single study of Read coding in primary care data alone reported a PPV of 81%, increasing to 90% with the presence of a relevant medication code in addition to a diagnostic code[20]. The two studies of medication data alone reported PPVs of 53% and 87%[17,21]. The single, small study of mortality data had a PPV of 67%[12].

**Fig 2:**
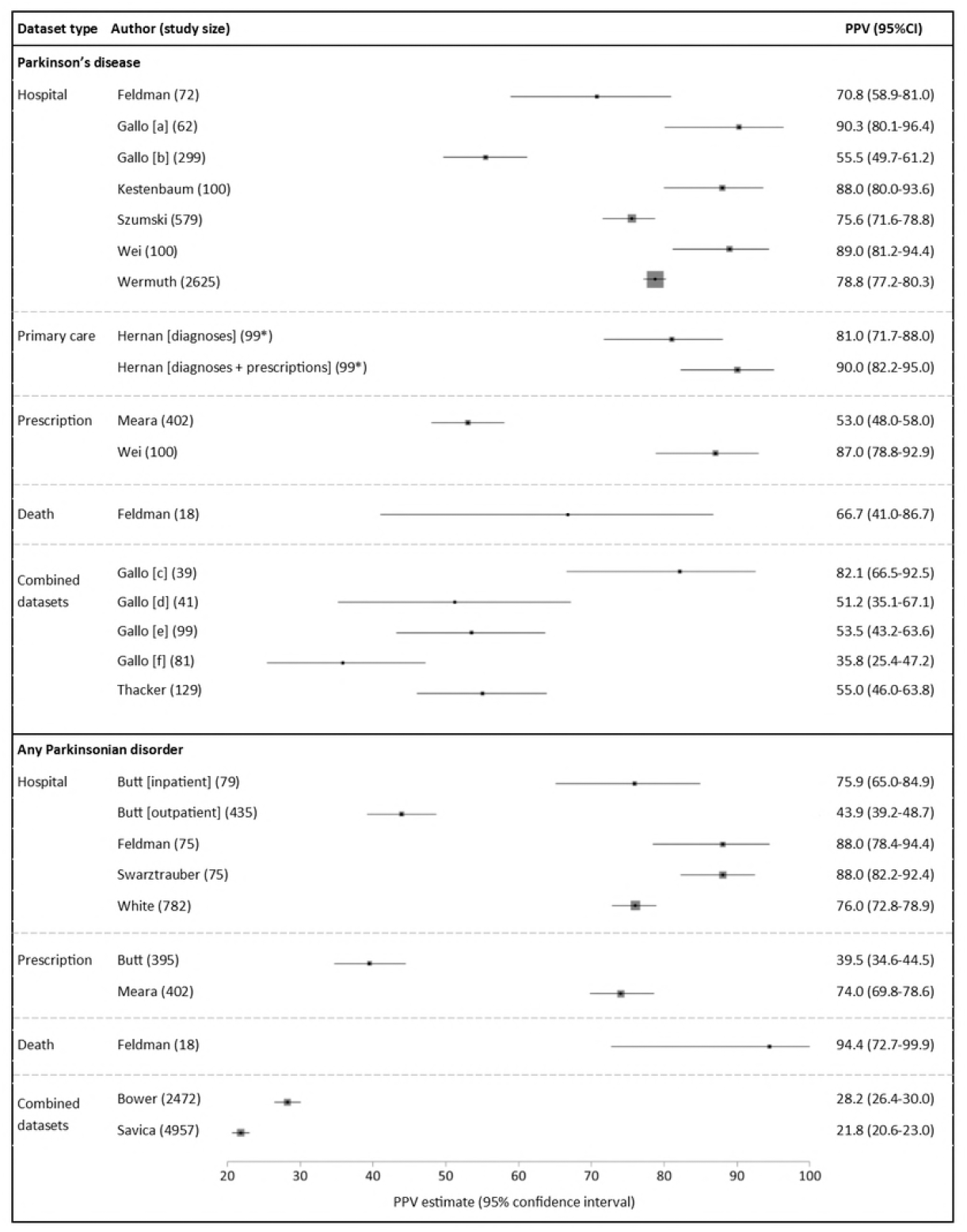
Positive predictive values (PPVs) of coded diagnoses. Study size: total number of code-positive cases (true positives + false positives). *Exact sample size unknown, most conservative estimate used. Box sizes reflect Mantel-Haenszel weight of study (inverse variance, fixed effects).

Several within-study comparisons were available from three studies identifying PD (Table 3)[12,16,17]. Two of these investigated the change in PPV for hospital data to identify PD when algorithms containing additional criteria were used[12,16]. Both showed a moderate increase in PPV if a relevant diagnosis code was recorded more than once, or if a specialist department assigned such a code. One study reported an increase in PPV when only primary position diagnoses were assessed[12]. Another showed that incorporating selected medication codes with diagnosis codes increased the PPV from 76% to 86%, although this was at the expense of reduced case ascertainment[16]. Finally, one study showed that the combination of a diagnostic code in hospital data with a relevant medication code increased the PPV when compared to using either dataset alone (94% versus 87% and 89% respectively)[17].

For parkinsonism there were 10 PPV estimates in total (Fig 2)[11,12,15,19,21–23]. These comprised five estimates from hospital data alone[11,12,15,19], two from prescription data alone[11,21], one from mortality data alone[12], and two from using datasets in combination[22,23]. PPVs ranged from 40-94% in the single datasets and from 22-28% in the combination datasets. The two studies of parkinsonism in prescription data produced very different PPV estimates of 40% and 74%[11,21]. One of these studies reported that the PPV of medication data to identify any parkinsonian disorder was considerably higher than that for PD (74% and 53% respectively)[21].

**Table 3:**
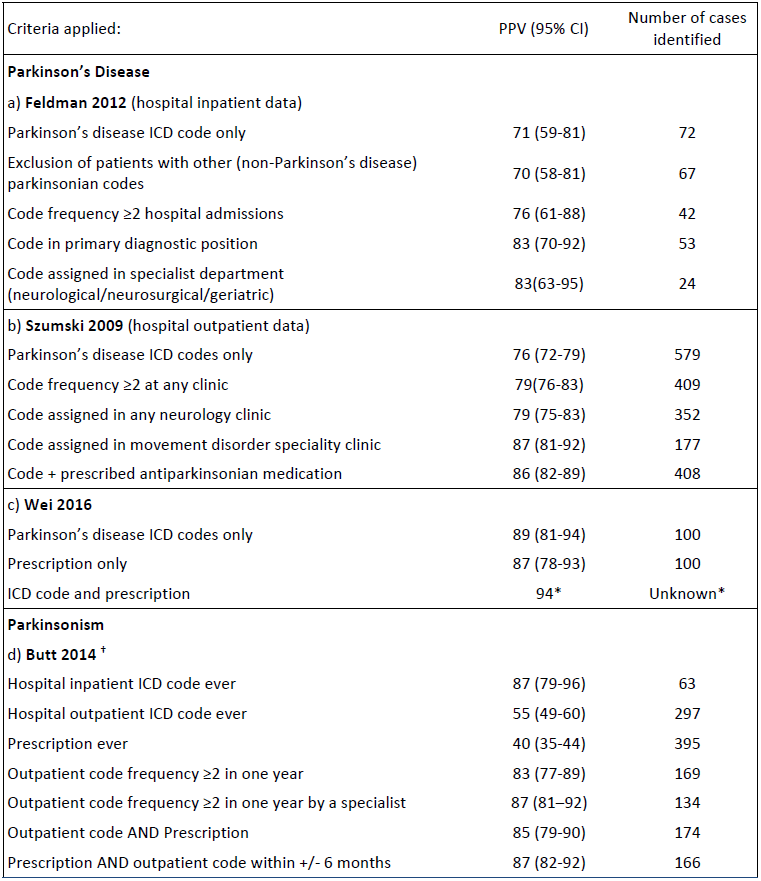
Within-study analyses: algorithm development.

| Criteria applied: | PPV (95% CI) | Number of cases identified |
| --- | --- | --- |
| **Parkinson's Disease** |  |  |
| **a) Feldman 2012** (hospital inpatient data) |  |  |
| Parkinson's disease ICD code only | 71 (59-81) | 72 |
| Exclusion of patients with other (non-Parkinson's disease) parkinsonian codes | 70 (58-81) | 67 |
| Code frequency $\geq 2$ hospital admissions | 76 (61-88) | 42 |
| Code in primary diagnostic position | 83 (70-92) | 53 |
| Code assigned in specialist department (neurological/neurosurgical/geriatric) | 83(63-95) | 24 |
| **b) Szumski 2009** (hospital outpatient data) |  |  |
| Parkinson's disease ICD codes only | 76 (72-79) | 579 |
| Code frequency $\geq 2$ at any clinic | 79(76-83) | 409 |
| Code assigned in any neurology clinic | 79 (75-83) | 352 |
| Code assigned in movement disorder speciality clinic | 87 (81-92) | 177 |
| Code + prescribed antiparkinsonian medication | 86 (82-89) | 408 |
| **c) Wei 2016** |  |  |
| Parkinson's disease ICD codes only | 89 (81-94) | 100 |
| Prescription only | 87 (78-93) | 100 |
| ICD code and prescription | 94* | Unknown* |
| **Parkinsonism** |  |  |
| **d) Butt 2014 <sup>†</sup>** |  |  |
| Hospital inpatient ICD code ever | 87 (79-96) | 63 |
| Hospital outpatient ICD code ever | 55 (49-60) | 297 |
| Prescription ever | 40 (35-44) | 395 |
| Outpatient code frequency $\geq 2$ in one year | 83 (77-89) | 169 |
| Outpatient code frequency $\geq 2$ in one year by a specialist | 87 (81-92) | 134 |
| Outpatient code AND Prescription | 85 (79-90) | 174 |
| Prescription AND outpatient code within +/- 6 months | 87 (82-92) | 166 |

The effect of additional criteria to identify PD cases on PPV and the number of cases identified. * Sample size and confidence intervals unknown for this accuracy measure.

### Sensitivity

For PD, there were 11 sensitivity estimates in total (Fig 3)[12,25–28]. Of these, nine were sensitivity estimates for mortality data alone, consistently showing that codes in the primary position only gave low sensitivities of 11-23%, rising to 53-60% when codes from any position were included[12,25–28]. A single study reported the sensitivity of hospital data to be 73%, increasing to 83% when hospital and mortality data were combined. There were no sensitivity estimates for primary care or prescription data.

For parkinsonism, there were three sensitivity estimates, all from one study[12]. Hospital admissions and mortality data combined gave higher sensitivity (71%) compared with either mortality or hospital data alone (43% and 63% respectively).

**Fig 3:**
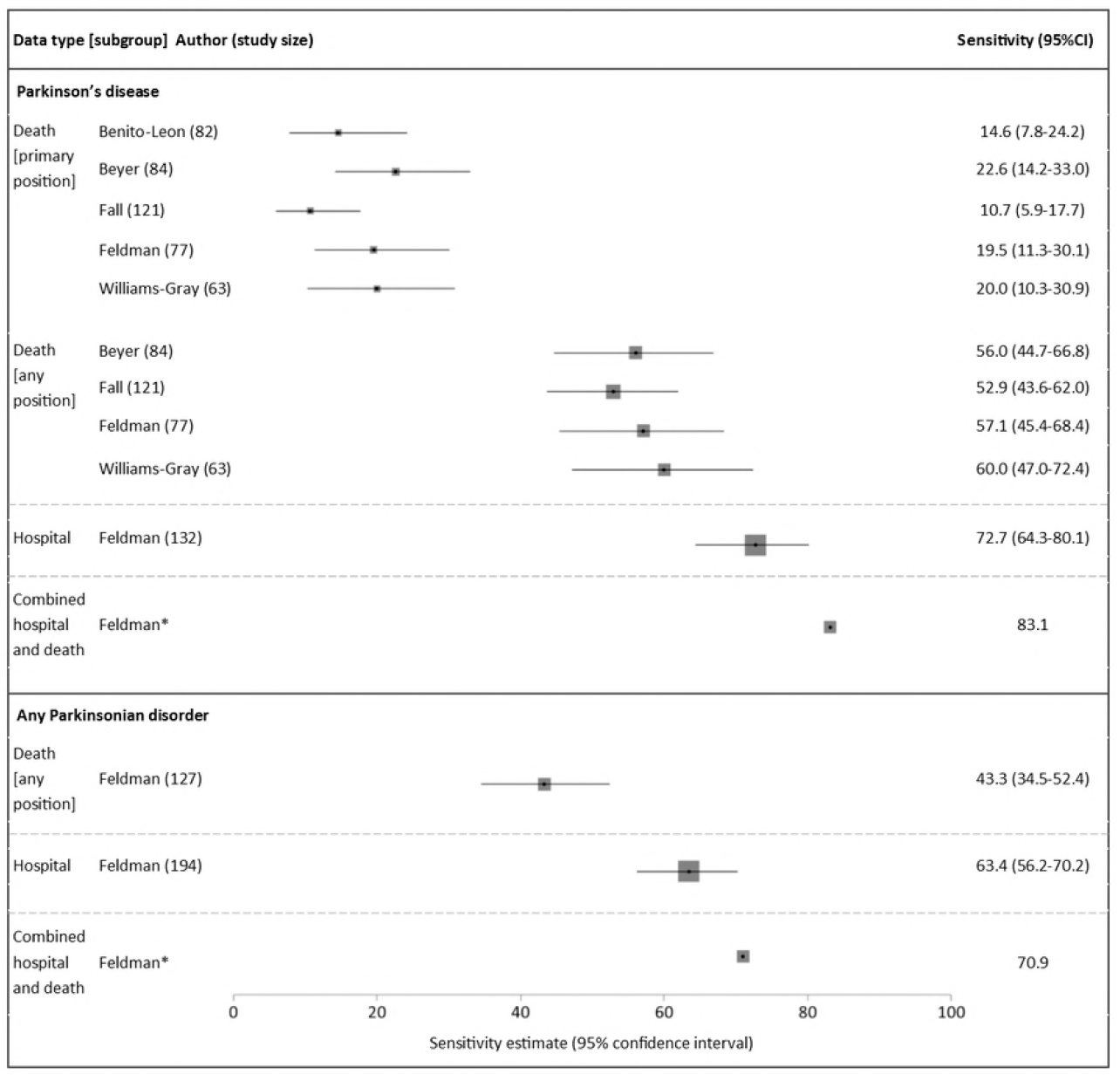
Sensitivity estimates of coded diagnoses. Study size: total number of true positives according to reference standard (true positives + false negatives). *Unknown sample size and confidence intervals. Box sizes reflect Mantel-Haenszel weight of study (inverse variance, fixed effects).

## Discussion

We have demonstrated that existing validation studies show a wide variation in the accuracy of routinely-collected healthcare data for the identification of PD and parkinsonism cases. Despite this, in the right setting, achieving high PPVs is possible. Sensitivity is generally lower than PPV, but is increased by combining data sources.

False positives (participants who receive a disease code but do not have the disorder) may arise in routinely-collected coded datasets for several reasons. Firstly, the clinician may incorrectly diagnose the condition. Given that PD and other parkinsonian disorders are largely clinical diagnoses made without a definitive diagnostic test, there is the potential for diagnostic inaccuracies. Clinicopathological studies have shown discrepancies between clinical diagnoses in life and neuropathological confirmation[29] and there is evidence that accuracy increases when diagnoses are made by movement disorder specialists[30–32]. Secondly, diagnoses may be incorrectly recorded in medical records, or errors may arise during the coding process. Similarly, false negatives (patients who have the condition but do not receive a code) may arise due to under-diagnosis, omission of the diagnosis from the medical records (e.g., because the condition is not the primary reason for hospital admission), or errors during the coding process.

The pharmacological treatment of PD is largely focussed on improving motor function and patients are treated with a limited number of drugs. This has allowed antiparkinsonian drugs to be used as ‘tracers’ in epidemiological studies[33,34]. There are potential problems with using prescription data as a proxy for PD diagnosis. This approach may disproportionately under-identify patients with early stage disease who do not yet require treatment. Also, a response to a trial of dopaminergic drugs may be used as part of the diagnostic assessment in potential PD cases, meaning some patients prescribed antiparkinsonian medications will not be subsequently diagnosed with PD. Furthermore, antiparkinsonian can be prescribed for indications other than PD (such as dopamine agonists for restless legs syndrome, endocrine disorders and other forms of parkinsonism). The specific drugs licensed for use in parkinsonian conditions varies between countries and may change over time. Therefore, an algorithm incorporating prescription data would need to be continually revised to match prescribing patterns. Results from our review suggest that prescription data alone has a low PPV for PD case ascertainment[21]; however, when drug codes are combined with diagnostic codes, PPV increases but with reduced case ascertainment[16,20]. Furthermore, prescription datasets appear to have a higher PPV when identifying any parkinsonian disorder rather than specifically PD[21].

This study has several strengths and limitations. Our review benefits from prospective protocol publication, comprehensive search criteria, and independent duplication of each stage by two authors. Despite this, relevant studies may still have been missed, especially if a validation study was a subsection of a paper with a wider aim. As all eligible studies were included, the results may have been influenced by studies of lower quality. Only two articles were found to be at low risk of bias or applicability concerns[11,12], and it is likely that biases in study design would have affected the results. For example, one study with the lowest PPV[23] used very broad ICD-9 codes such as 781.0 (abnormal involuntary movements) and 781.3 (lack of coordination).

Since there is no method of diagnosing PD with certainty in life, there is likely to be some misclassification of the reference standards used in the studies. The application of stringent diagnostic criteria to reference standard diagnoses, although often necessary for research purposes, may lead to some patients being misclassified as ‘false positives’ when they do in fact have the condition. This may lead to underestimation of the PPV in some of the studies. When considering the ideal reference standard for validation studies, there is a trade-off between the robustness of the reference standard and validating sufficient cases to produce precise accuracy estimates. For example, in-person neurological examination may have greater diagnostic certainty than medical record review but this becomes difficult as the cohort size increases.

Many of the studies reported cases with insufficient information to meet the reference standard and the handling of these varied. Some studies excluded such cases, others classified them as false positives, while some did not specify how they handled such missing data. Excluding such cases may introduce selection bias, whereas counting them as false positives may underestimate PPV.

The effect of possible publication bias on the results is difficult to estimate, but disproportionate publication of studies which report more favourable accuracy measures may lead to over-estimation of the performance of the codes. In addition, estimates of PPV are dependent upon the prevalence of the condition in the study population but it was not possible to assess the prevalence of PD within each study population.

Our review highlights several areas requiring further research. Given that the management of PD is largely delivered in outpatients or the community, primary care data may be an effective method of identifying cases. Whilst studies have suggested that PD diagnoses made in primary care are less accurate than those made in a specialist setting[35,36], primary care records combine notes made by primary care clinicians with prescription records and correspondence from secondary care. Codes from primary care should therefore include diagnoses made by specialists, thus increasing their accuracy. We found only one small study of primary care data, reporting a promising PPV of 81%, improving to 90% with the inclusion of medication codes[20]. No studies investigated the sensitivity of primary care data. Further research into the accuracy of primary care data is needed.

Two studies investigated using algorithmic combinations of codes from different sources to improve PPV[12,16]. These investigated the additional benefit of the inclusion of factors such as only including codes that appeared more than once, selecting codes in the primary position only, combining diagnostic codes with prescription data, and only including diagnoses made in specialist clinics. These methods increased PPV but at a cost to the number of cases identified. The development of algorithms that maximize PPV whilst maintaining a reasonable sensitivity (e.g., by combining multiple complimentary datasets) merits further evaluation.

To our knowledge, no studies have evaluated the accuracy of routinely-collected healthcare data for solely identifying atypical parkinsonian syndromes such as PSP and MSA. Further work is needed to understand whether these datasets provide a valuable resource for studying these less common diseases.

In conclusion, our review summarises existing knowledge of the accuracy of routinely-collected healthcare data for identifying PD and parkinsonism, and highlights approaches to increase accuracy and areas where further research is required. Given the wide range of results observed, prospective cohorts may wish to perform their own validation studies based on their specific setting and research question.

## Acknowledgements

The UK Biobank Neurodegenerative Outcomes Working Group provided feedback on the manuscript. This work was conducted on behalf of Dementias Platform UK (www.dementiasplatform.uk/)

## Supporting information

S1 File. Search strategy.

S2 File. QUADAS-2 assessment.

S3 Table. QUADAS-2 summary results.

S4 Checklist. PRISMA checklist.

